# Active mucus-cilia hydrodynamic coupling drives self-organisation of human bronchial epithelium

**DOI:** 10.1101/2019.12.16.878108

**Authors:** E. Loiseau, S. Gsell, A. Nommick, C. Jomard, D. Gras, P. Chanez, U. D’Ortona, L. Kodjabachian, J. Favier, A. Viallat

## Abstract

The respiratory tract is protected by mucus, a complex fluid transported along the epithelial surface by the coordinated beating of millions of microscopic cilia, hence the name of mucociliary clearance. Its impairment is a strong marker of severe chronic respiratory diseases. Yet, the relationship between ciliary density and the spatial scale of mucus transport, as well as the mechanisms that drive ciliary-beat orientations during ciliogenesis are much debated. Here, we show on polarized human bronchial epithelia that mucus swirls and circular orientational order of the underlying ciliary beats emerge and grow during ciliogenesis, until a macroscopic mucus transport is achieved for physiological ciliary densities. By establishing that the macroscopic ciliary-beat order is lost and recovered by removing and adding mucus respectively, we demonstrate that cilia/mucus hydrodynamic interactions govern the collective dynamics of ciliary-beat directions. We propose a two-dimensional model that predicts a phase diagram of mucus transport in accordance with the experiments. It paves the way to a predictive in-silico modeling of bronchial mucus transport in health and disease.

## Introduction

The airways are protected from inhaled pollutants and pathogens by a layer of mucus, a complex fluid that is continuously transported along the bronchial walls before being cleared out and swallowed. The mucus is propelled via the continuous beating of active cilia exposed by the epithelial ciliated cells. This process, called mucociliary clearance, is strongly impaired in chronic respiratory diseases, such as primary dyskinesia, severe asthma^1^, COPD ^2^ and cystic fibrosis ^3^, which affect hundreds of millions of people^4^.

Effective directional transport of mucus requires both a high surface density of active epithelial cilia to overcome friction forces on the epithelial wall and a directional coordination of ciliary beats. Recently, centimetric mucus swirls and a circular orientational order of the underlying ciliary beats ^5^ were observed in bronchial epithelia reconstructed in vitro only when the ciliary density was high ^5,6^. Yet, the relationship between cilia density and the spatial scale on which mucus is transported is unknown. Furthermore, the biological and physical mechanisms involved in the emergence of the orientational order of ciliary beats during ciliogenesis, its stability and long-term repair are much debated. The ciliary beat direction is set by the polarity of the basal body of the cilium and is considered to be fixed during development by the planar cell polarity (PCP) proteins that regulate the polarization of epithelial tissue ^7–9^. However, daily variations of ciliary-driven cerebrospinal fluid flow patterns, recently reported in epithelia of mouse brain ventricles, showed that ciliary beats can change direction ^10^. Moreover, under an external fluid flow, the ciliary beat directions on the skin of the Xenopus embryo before PCP is in place and on maturing cell cultures of mouse ependyma redirected along the flow streamlines ^11,12^. Cilia can then actively respond to external hydrodynamic cues. Interestingly, the bronchial epithelium is naturally continuously subjected to the endogenous mucus flow generated by ciliary beats. This raises crucial questions: could mucus flow affect the long-range ordering of ciliary beats by creating a hydrodynamic coupling among cilia? Would hydrodynamic feedback on cilia lead to an enhancement of orientational ciliary beat order, and in turn result in a better mucus transport?

Here, we address these questions by combining experiments on in-vitro reconstituted human bronchial epithelia and a hydrodynamic model. We observe and quantify the dynamics of formation and growth of swirly patterns of mucus flow during ciliogenesis. We show that, at a physiological cilia density, a macroscopic swirl of mucus develops associated with a circular coordination of ciliary beat directions and a circular pattern of planar cell polarity of the tissue. By establishing that the macroscopic ciliary-beat order is lost and recovered by respectively removing and adding mucus, we demonstrate that cilia/mucus hydrodynamic interactions drive the collective dynamics of ciliary-beat directions. Based on these observations, we developed a two-dimensional hydrodynamic model which highlights two relevant physical parameters of the epithelium, the density of cilia and mucus viscosity, allowing the establishment of a phase diagram of mucus transport.

## Results

The reconstituted bronchial epithelium is cultured in-vitro at the air-liquid interface from human primary cells (Fig. 1a). The pseudo-stratified tissue consists of basal cells, goblet cells and ciliated cells each carrying 200-300 active cilia gathered in bundles on the apical surface ^13^. After cellular differentiation, the tissue exhibits the characteristics of the bronchial epithelium ^14^ such as active cilia beating at a frequency of about 8-10 Hz and the production and transport of mucus on the epithelium surface. In the culture chamber, the basal side is in contact with the culture medium through a porous membrane and the apical part is at the air interface. Mucus flows and beats of bundles of cilia are measured by videomicroscopy.

**Figure 1.**
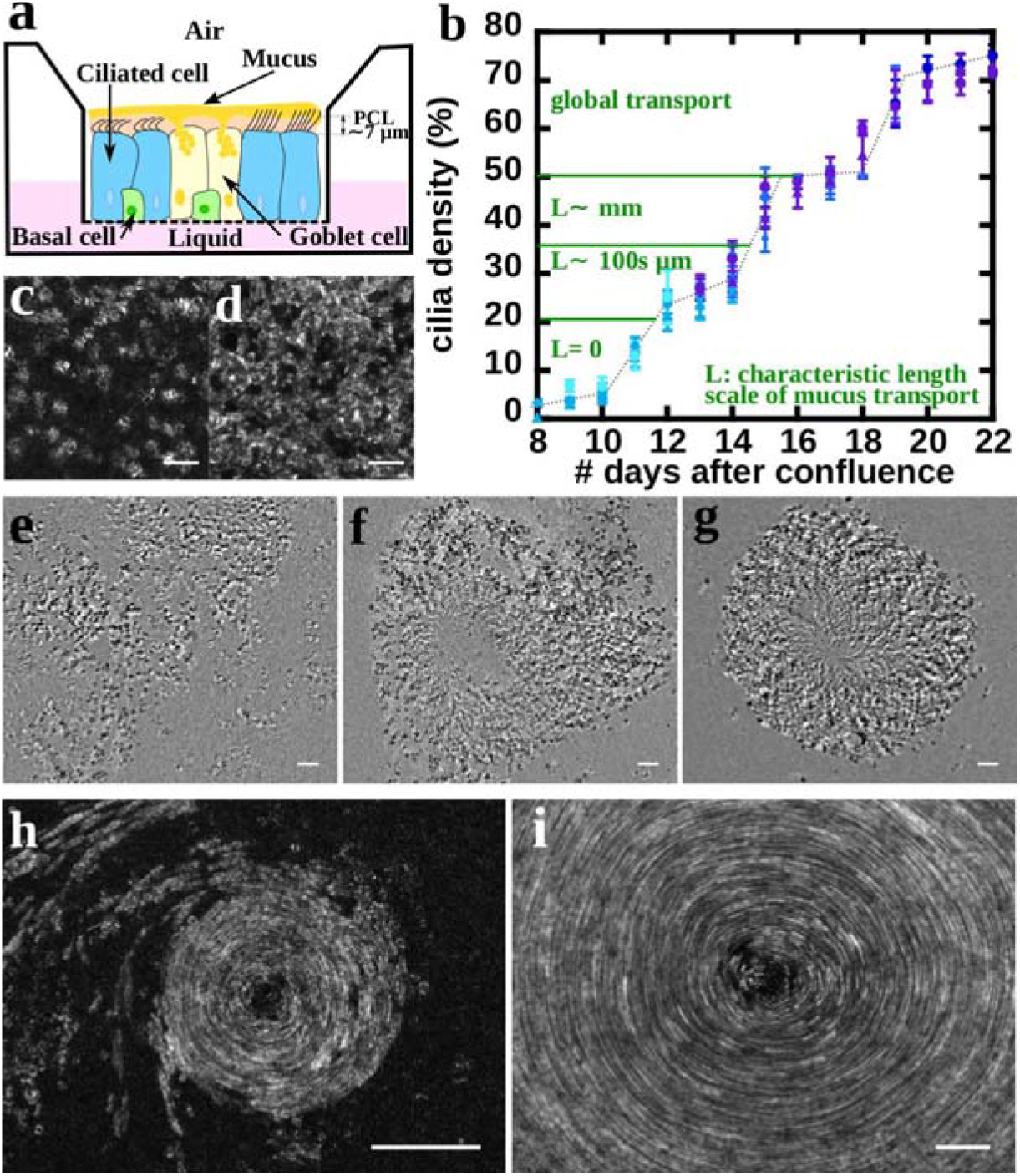
Emergence of mucus flow patterns during ciliogenesis. **a**, Schematic of the ALI chamber: the basal side of the bronchial epithelium is in contact with the culture medium through a porous membrane and the apical side points towards the air. The epithelium has a pseudo stratified structure made of basal cells, goblet cells, which produce the mucus and ciliated cells. During the forward stroke, cilia tips poke in the mucus layer while their recovery motion takes place in the periciliary layer. **b**, Dynamics of the ciliogenesis. The curve represents the temporal evolution of the percentage of surface covered by active cilia after initiation the differentiation. Data are from samples #1 to #10 from 5 patients, the color represents a patient and different symbols of the same colors represent different cultures from the same patient. Each data point is an average over at least an area of 1mm × 1mm and error bars are the standard deviation. Characteristic length scale of mucus transport associated to the cilia density are written in green. **c-d**, Images representative of two cilia densities, *φ* = *30%* in **c** and *φ* = *80*% in **d**. Images are the result of a standard deviation projection over 40 frames acquired at 40 fps. Scale bars = 30 μm. **e-g**, Formation of a swirl of mucus from day 12 to day 14 in sample #13. The three images are the same field of view. The background has been subtracted to better visualize the debris embedded in the mucus by subtracting to each frame of the movies S2-S4 the median of the temporal projection. Scale bars = 50 μm. **h-i**, growth of a mucus swirl between day 12 and day 15, in sample #8, by successive accumulation of mucus transported in the vicinity of the swirl. Images results from the standard deviation projection over 100 frames at 0.2 fps in **h** and over 150 frames at 40 fps in **i**. Scale bars = 100 μm.

### Ciliogenesis dynamics: emergence and growth of mucus swirls and ciliary-beat coordination

#### Density of active cilia

The percentage of the culture area covered by the active cilia, φ, was measured daily from the full confluency of the cell culture and throughout ciliogenesis on 10 cultures from 5 donors (see Fig. 1b and Table S1). After eight days, φ is lower than 5% (Fig. 1b). Then, the ciliary density increases by successive steps of 20 to 25% (Fig. 1c) to reach a final value of about 70-80% after two more weeks (Fig. 1d). Interestingly, the dynamics of ciliogenesis is almost identical for all 10 cultures (Fig. 1b).

#### Spatial range of mucus transport

From day 8 to day 11, when φ increases from 3% to 15%, the mucus layer above the cilia vibrates but is not transported (see Movie S1). A local mucus transport is observed on day 12 (Fig. 1e and Movie S2), for φ ~ 25%. Strikingly, from day 13 and 14, while φ remains almost constant, mucus swirls of a few hundred micrometers in diameter (Fig. 1f, g and Movies S3 and S4) are observed, together with small paths along which mucus is transported towards the edges of the culture. When φ gradually increases, the number of swirls decreases but their diameter increases by capturing the mucus transported nearby (Fig. 1 h & i, Movie S5) and reaches the millimeter scale for φ in the 35-50% range. For φ greater than 50%, a single mucus swirl extends over the entire surface of the culture (Fig. 1i and Movie S6). The characteristic size of mucus swirls during ciliogenesis is indicated in Fig. 1b.

#### Orientational order of the ciliary beats

the beat direction of each bundle of cilia is represented by a rod in Fig. 2a and 2b. As illustrated in Fig. 2b, the directions of the ciliary beats present a strong circular order under the mucus swirls, with only a few localized defects mainly located in the swirl center. Areas where ciliary beats are aligned are also observed (Fig.2a). The beat orientational order is quantified in a square surface of length L by 2 parameters classically used for nematic materials ^15,16^. The first one is a local order parameter S(L) = <cos 2θ>^2^ + <sin 2θ>^2^, where θ is the angle between the beat direction and a reference direction, and the average < > relates to all bundles in the domain of size L. S varies between 0 and 1; 0 and 1 are respectively an isotropic state with random beat directions and a fully ordered state with aligned beat directions. The second parameter is the density of topological defects defined as local discontinuities in the director field of beat directions (Fig. 2a). They are associated with topological charges + ½ (Fig. 2a red circles) or −½ (Fig. 2a blue triangles), corresponding to a clockwise or counterclockwise π-rotation of the director field around the singularity respectively.

**Figure 2.**
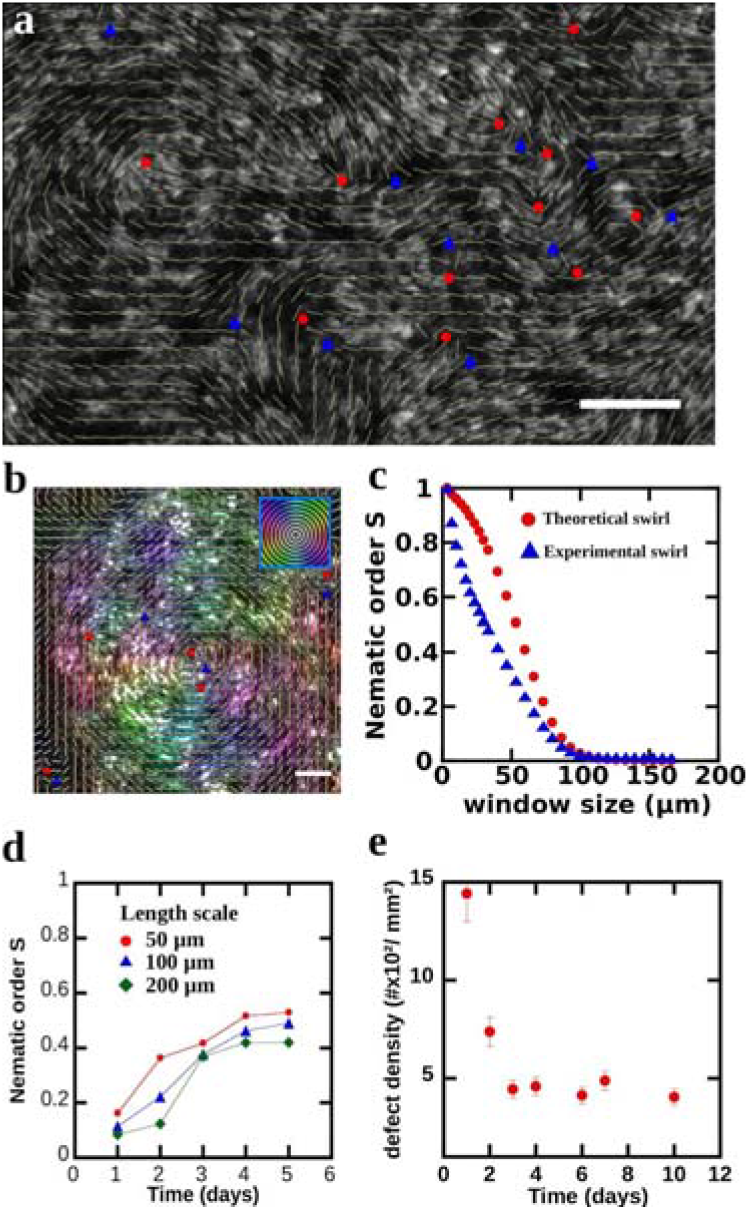
Orientational order of ciliary beats. **a,** the beating directions of cilia is revealed by performing a standard deviation projection over 40 frames at 40 fps. The director field of the beating directions is plotted as an overlay and is computed by averaging the orientation in boxes of 15×15 pixels, corresponding to the size of a single ciliated cell. Red circles and blue triangles are located where a discontinuity arises in the director field and are +½ and −½ nematic topological defects respectively. A schematic of +½ and −½ defect is drawn in inset. Scale bar = 30 μm. **b**, Beating directions of cilia underneath a local swirl of mucus, sample #11. The colors code for the direction of beating according to the orientation scale in the upper right corner. The director field averaged over 15×15 pixels reveals a robust circular orientational order. Only few topological defects are present. Scale bar = 20 μm. **c**, Nematic order S, blue curve, associated to the cilia beating orientations of the panel **b**. S is computed in a sliding window of size L with L varying from L_min_= 6 μm, the size of a ciliated cell to L_max_ = 170 μm, the size of the swirl. The window is moved over the entire surface of the image by L/3 steps. The comparison of the nematic order of the cilia, blue curve, with the nematic order computed for a perfect circular orientation, red curve, shows that underneath a swirl of mucus, the cilia exhibit a strong circular order. **d**, Dynamics of ordering of cilia in a sample exhibiting a single swirl spanning the entire culture. The parameter order S is measured at mid-distance (~1.5 mm) between the center of rotation and the border of the chamber and is averaged on a total area of ~1mm^2^. Day 1 defines the first day the swirl spans the culture. At a length scale of 50 μm we observe a sharp increase of order within 24h, this indicates that cilia get more aligned overtime. At a larger length scale of 200 μm there is a 24h delay time before we observe the same increase of S. After four days, S reaches a plateau at ~0.5. **e**, Temporal evolution of the density of topological defects in the same experimental conditions than in **d**. Underneath the rotation of mucus, the number of defect decreases within a couple of days and then remains constant.

The ciliary-beat orientational order under a mucus swirl is computed at different length scales L. S(L) is averaged on a set of square tiles of length L that totally pave the mucus swirl, and is compared to S(L) computed for a full order when all ciliary-beat directions are tangential to the swirl. The comparison of the two curves shown in Fig. 2c for a typical ciliary beat 100 μm-vortex reveals a strong tangential orientational order of the ciliary beats.

At the end of ciliogenesis, below the macroscopic mucus swirl, we observed in its central part a strong circular orientational order of ciliary beats originating from a previous local pattern. Yet, farther from the center, at a distance of 1.5 mm, the ciliary-beat orientational order is low (S~ 0.1) (Fig. 2d). This order measured at different length scales (L= 50, 100 and 200 μm) increases very significantly within four days before reaching a plateau where S ranges between 0.4 to 0.55 (Fig. 2d). Simultaneously, the surface density of topological defects decreases threefolds (Fig. 2e). These results clearly show that the ciliary-beat directions continue to coordinate for several days on mature cultures. It should be noted that during this increase in order, mucus rotates on the epithelial surface where it exerts a continuous hydrodynamic stress on the top of cilia.

##### Planar cell polarity of the tissue

We performed immunofluorescent staining to image the core PCP protein Vangl1 to evaluate whether the cultures were polarized and to measure the main directions of planar cell polarity. We observed large domains where Vangl1 was aligned (fig. 3a-c, typical scale 200 μm) or circularly organised (Fig. 3d-f, typical scale 100 μm). These domains likely correspond to areas of linear mucus transport towards the culture edges and to local mucus swirls respectively. At a millimetric scale, a circular order of Vangl1 is also observed under an extended millimetric mucus swirl, as illustrated in Fig. 3g. In summary, the characteristic size of mucus transport estimated by the size of the spontaneously emerging mucus swirls increases strongly with cilia density to reach the entire culture size at physiological cilia density. These swirls are associated with both a circular order of ciliary-beat directions and a circular planar cell polarity of the tissue below the swirls. To understand the specific role of the hydrodynamic forces exerted by mucus on the coordination of ciliary beat directions, we removed and added mucus on mature epithelial surfaces.

**Figure 3.**
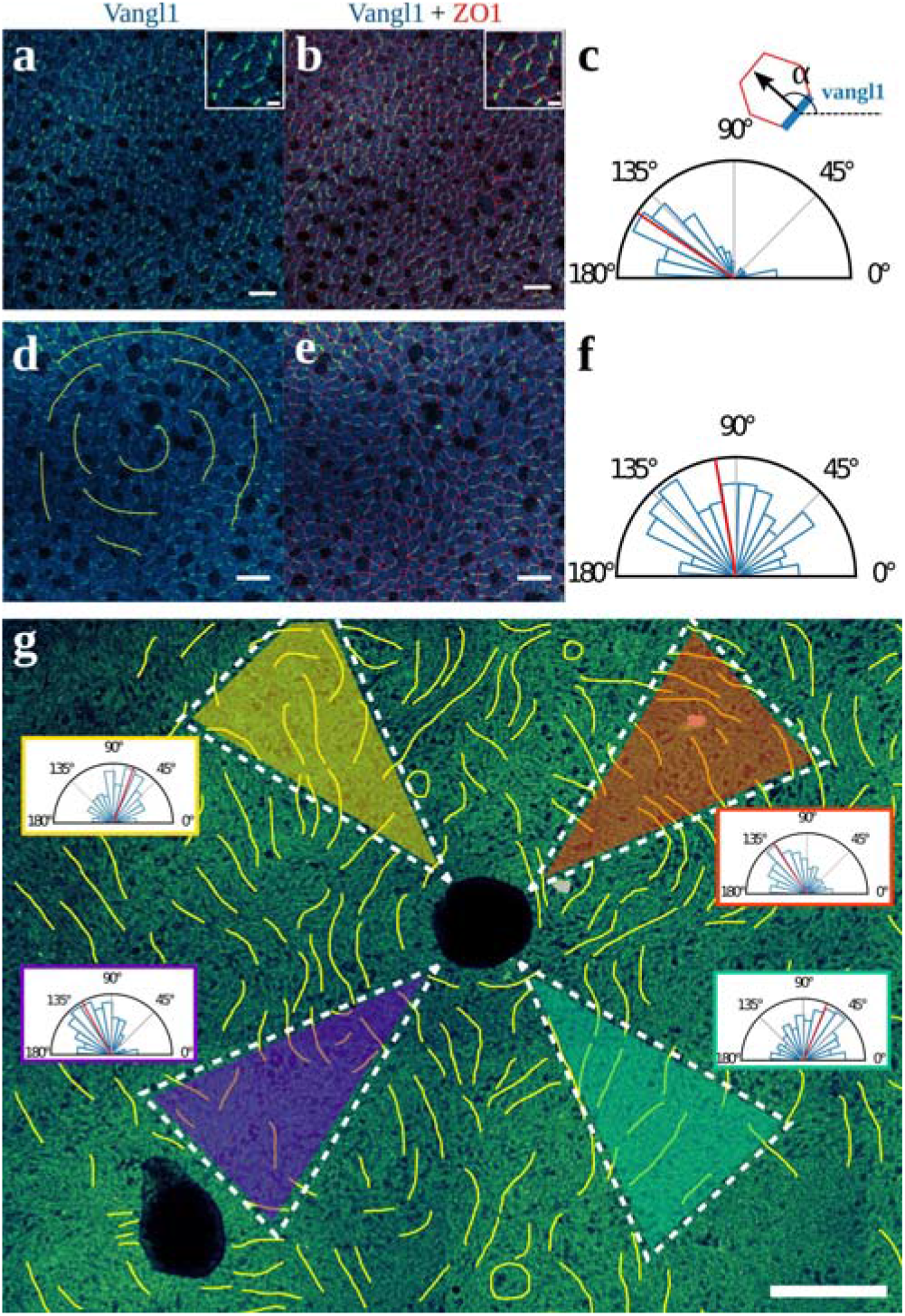
Planar cell polarity organisation. **a & b,** Long range linear polarization of the PCP Vangl1 protein (blue-green label). Tight junctions, ZO1, are labelled in red. Scale bars = 20 μm. In insets, a zoom on a few cells is shown. Scale bars = 5 μm. **c,** Quantification of the direction of ciliary beats according to the orientation of Vangl1: the beat direction is orthogonal to the Vangl1 edge. The angle distribution of beat directions corresponds to the field in **a** & **b,** the red line indicates the median direction of 146° and the standard deviation is 34°. **d & e,** Vangl1 exhibits a circular pattern reminiscent of the observed local swirls of ciliary beats. The yellow lines are a guide for the eyes and represent the main orientations orthogonal to vangl1. Scale bars = 20 μm. **f,** Angle distribution of beat directions, computed from Vangl1 orientation, corresponding to the field in **d** & **e**, angles are well distributed over the 0-180° and the red line indicates the median direction of 99°. A perfect circular order would lead to a median of 90°. **g,** PCP protein Vangl1 over an area of ~1.5 mm × 1 mm in a sample where a millimetric mucus swirl was present. Black areas are due to undulations of the tissue made during mounting of the sample. The yellow lines are a guide for the eyes and represent the main orientations orthogonal to Vangl1. They show the existence of a circular orientational order at the millimetric scale. Angle distributions of ciliary beat, computed from Vangl1 orientation, are plotted for 4 quadrants (dashed triangles, arc of a circle with radius of 500 μm and subtending an angle of 35°). The red lines indicate the median of the distributions with α = 73 for top left, α = 123 for top right, α = 68 for bottom right and α = 119 for bottom left. Scale bar = 200 μm.

### Mucus drives the loss and recovery of the ciliary orientational order

We removed rotating mucus on the surface of mature cultures by an extensive apical washing. Such washing empties the goblet cells thus limiting the production of new mucus for a week. Three days after mucus removal, the giant ciliary-beat orientational order spanning the culture chamber was lost and several local circular patterns emerged instead (Fig. 4a). Many topological defects appeared at a high density of about 6.10^2^ to 8.10^2^ defect/mm^2^. S decreases to 0.1 over a length scale of L= 300 μm compared to S=0.4 before mucus removal (Fig. 4b). Strikingly, a similar loss of ciliary-beat orientational order was observed when mucus got dehydrated at top of the epithelium and was no more transported although cilia still beat below (Fig. 4b and movie S7). This shows that mucus transport is mandatory to maintain the macroscopic ciliary-beat orientational order.

**Figure 4.**
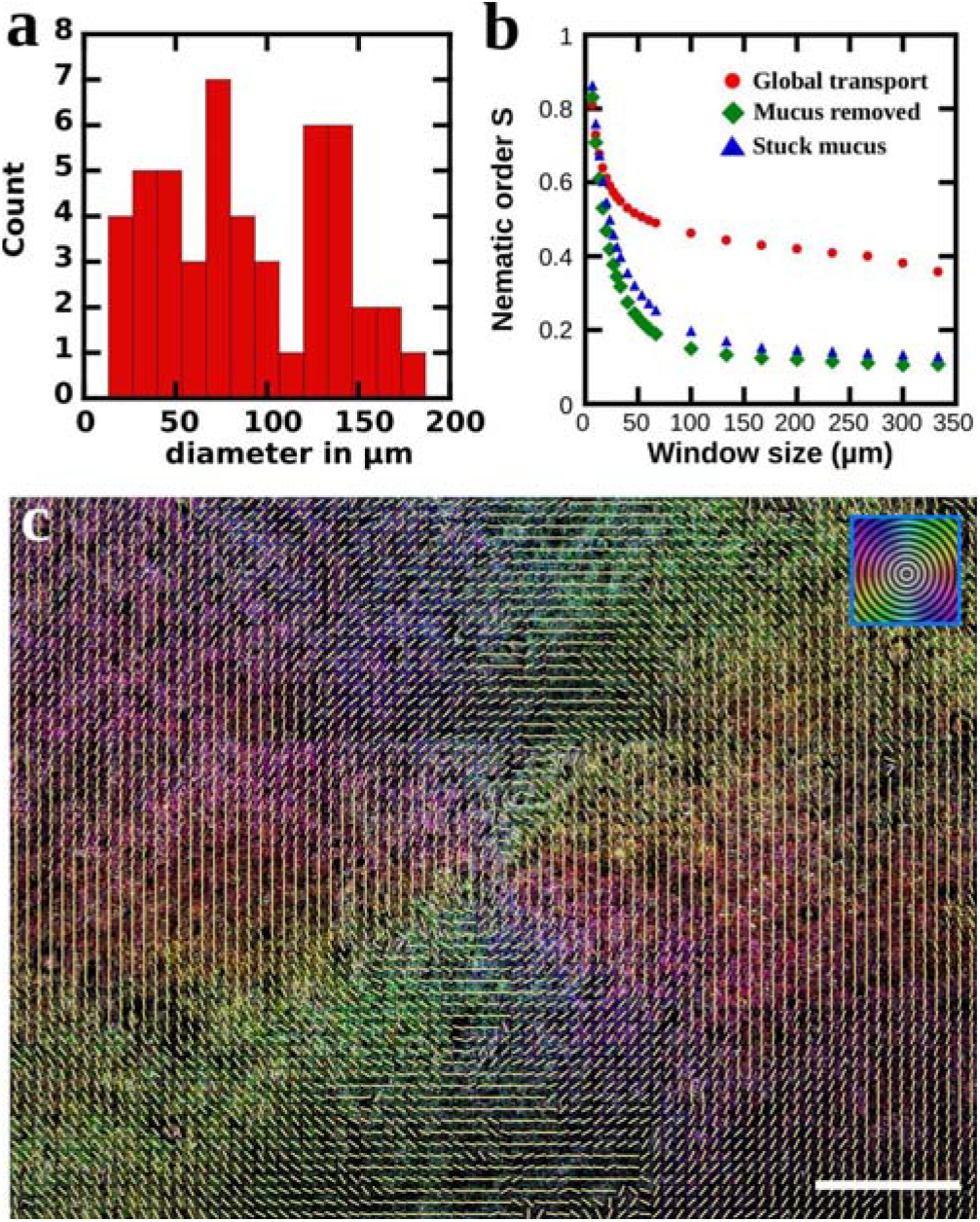
The transport of mucus drives the self-organisation of the epithelium. **a,** Size distribution of the local swirls that form after mucus removal. Data are pooled from samples #6, 10 and 11. **b,** Comparison of the nematic order computed at different length scales. When the mucus is removed or become immobile due to dehydration, we observe a loss of order at length scales greater than ~25 μm. **c**, Orientation map of the beating direction of cilia 3 days after addition of a diluted mucus on top of a disorganized culture (sample #15). The colors code for the beating direction according to the orientation scale in the inset. The director field is plotted by averaging the directions over a group of 10 cells. The bronchial epithelium exhibits a circular orientation of the ciliary beats over a millimetric distance. Scale bar = 200 μm.

Next, the mucus stored at the edge of the culture chamber was diluted and redistributed over the entire epithelial surface. After three days, a new millimetric mucus swirl was reconstituted at the epithelial surface (see movie S8), associated with a strong circular orientational order of underlying ciliary beats (Fig. 4c). The mucus flow forces ciliary beats to recover a large-scale orientational order. These results show that the direction of ciliary beats is not frozen after ciliogenesis in in-vitro reconstituted bronchial epithelium and supports the idea that cilia actively respond to the hydrodynamic force exerted by mucus.

To understand the underlying hydrodynamic mechanisms, we developed a simple hydrodynamic 2D-model.

### The hydrodynamic coupling between mucus and cilia drives the emergence of a global order

The modeling approach is based on the schematic view shown in Figure 5a. The epithelial surface is described using hexagonal unit elements of size D, which represent cells or groups of cells that can be ciliated or not. The fluid in contact with the epithelium is decomposed in two layers ^17^: the low-viscosity periciliary layer (PCL), where ciliary beating mostly occurs, and the high-viscosity mucus layer. We focus on long-range flow phenomena developing in directions parallel to the epithelium, over length scales much larger than the typical fluid layer thickness. Therefore, the mucus flow is considered to be two-dimensional. In this configuration, the PCL behaves as a slip layer. This approach, illustrated by the schematized velocity profiles in Figure 5a, is consistent with prior results issued from three-dimensional numerical simulations ^18,19^.

**Figure 5.**
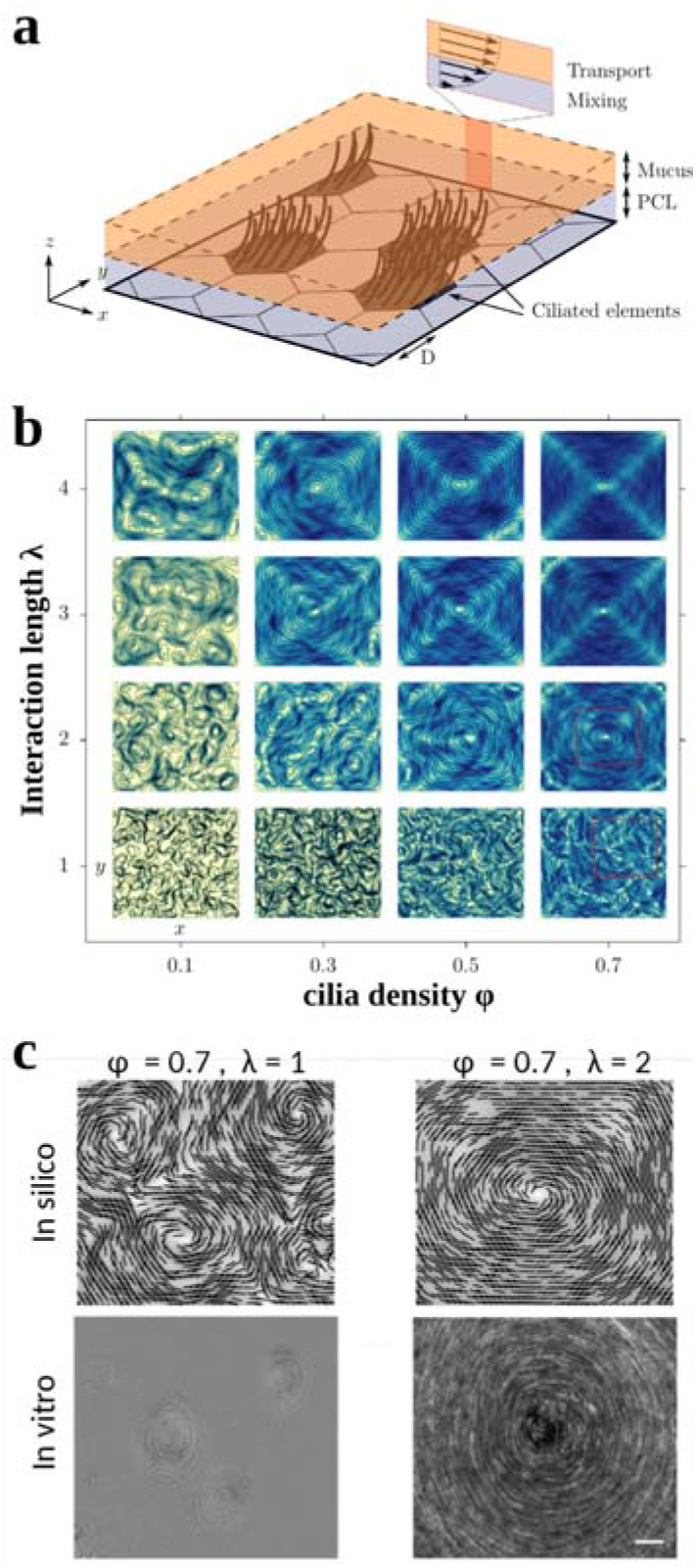
Hydrodynamic model of the bronchial epithelium. **a,** Schematic of the modelled epithelium. The 2-dimensional epithelium is paved with ciliated elements and non-ciliated elements of characteristic size D. The PCL is a shear region with the fluid velocity in the xy plane equal to zero on the epithelial surface and is equal to the mucus velocity at the PCL/mucus interface. The mucus is transported in the xy plane. **b**, Phase diagram of the mucus flow patterns in the λ, φ space. Panels show streamlines and iso-contours of the non-dimensional flow velocity magnitude. In all cases, the solution has reached a steady state, and the motile force on motile elements is aligned with the streamlines. At a given λ, increasing φ from 0.1 to 0.7 results in a transition from a local and small swirly patterns to a large global swirl. At a given φ, an increase of the hydrodynamic interaction length results in an increase of the characteristic size of the mucus flow patterns. Red rectangle indicates the part of the computational domain examined in c. **c**, Comparison between simulation and experiments. At a fixed cilia density of 0.7, typical of a mature culture, increasing λ from 1 to 2 results in a transition from multiple swirls to a single global swirl. In-vitro we observe the same mucus flow patterns on mature cultures in absence or in presence or mucus, respectively.

In this model, ciliary activity is averaged over time and space. We consider only the global driving force that propels the mucus in a specific direction by averaging over time the force resulting from the stroke recovery cycle of beats. At the scale of one hexagonal element, we average the driving forces exerted by all the cilia of the considered element.

The flow is modeled by the incompressible two-dimensional Navier-Stokes equations. The effect of the epithelial wall on the flow is modeled through by a body force (i.e. momentum source/sink) **f**_**n**_ exerted on the mucus. Two types of momentum sources are considered. The friction force relates to the normal shear stress in the PCL. It applies over the entire epithelial plane and is assumed to linearly depend on the mucus velocity **U**, **f**_**v**_ = −κ**U**, with κ a friction parameter. The ciliary force **f_c_**, locally driving the mucus in a specific direction as a result of time-averaged ciliary activity, is applied on ciliated elements. Its magnitude is constant and its orientation is forced to gradually align with the local mucus velocity, as detailed in *Materials and methods*. This alignment rule provides a hydrodynamic interaction mechanism between ciliated elements. The total momentum source in the flow equations is expressed as 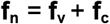

The model depends on three non-dimensional physical parameters, φ, the Reynolds number Re = ρD‖**f_c_**‖/κμ, with μ the mucus viscosity, and a non-dimensional hydrodynamic interaction length λ = (μ/κ)^1/2^/D. The latter parameter represents the typical range of influence of a ciliated element and defines the maximum interaction length between ciliated cells. The flow solution is computed over ranges of λ ∈ [0.1, 0.7] and φ ∈ [1, 4] in a square domain of ~7000 elements, using numerical simulations based on the Lattice-Boltzmann method, as detailed in Material and methods. The Reynolds number is set to 0.1. The initial location of ciliated elements and their initial orientation are randomly set. A free-slip boundary condition is used on the limits of the square domain in order to reproduce a confinement similar to that of the experiments.

We obtain a phase diagram which captures the observed mucus flow patterns (Fig. 5b). Indeed, mucus flows exhibit swirly patterns, whose typical length scale increases as a function of cilia density as experimentally observed during ciliogenesis. For φ ≥ 0.5 and λ ≥ 2, a global swirl spanning the entire chamber emerges, similar to what we observe for a cilia density of 50 % in presence of mucus. At a fixed cilia density of 70 %, the decrease of the interaction length λ from 2 to 1, equivalent to a decrease of the viscosity of the transported fluid, drives the transition from a global swirl to multiple local swirls. This λ dependence of the loss of the global orientational order is reminiscent of the observed transition which occurs when we remove the mucus and replace it with culture medium (Fig. 5c). The combination of the experiment with the hydrodynamical model provides a strong evidence that long range hydrodynamic interactions are sufficient to establish a global orientational order in the bronchial epithelium.

## Discussion

In this work we first described the fascinating spontaneous emergence of mucus swirls on reconstituted bronchial epithelia. We quantitatively measured the formation and growth of these swirls during ciliogenesis as well as the final long-range mucus transport observed for a cilia density higher than 50%. We highlight that the local milling patterns observed during ciliogenesis are not determined by the circular geometry of the culture chamber because they are numerous and randomly spatially distributed. However, the giant swirl observed at the end of ciliogenesis ‘feels’ the whole culture chamber and its circular shape is determined by the chamber geometry. Associated with these swirls, we showed that the ciliary beats and the planar cell polarity of the tissue have a circular orientational order.

We proposed a model that describes the mechanism of mucus swirl formation and coordination of ciliary-beat directions based on our observations that mucus is essential to maintain and recover large-scale transport. The model relies on the active coupled response of cilia whose beat direction locally tends to align with the mucus flow resulting from all surrounding ciliary beats. It predicts the emergence of mucus swirls through a circular orientation of ciliary beats, their growth with cilia density and with the length of hydrodynamic interactions, directly related to mucus viscosity. Noteworthy, the ciliary beat system belongs to the class of active nematic living materials, recently observed in several living organisms ^15,16,20^and whose studies aim to link nematic properties to biological functions. Here, the bronchial epithelium is an even more complex material composed of a double nematic system. Indeed, cilia are not only subjected to the mucus-driven hydrodynamic force, but are also constrained at their base by the PCP protein long-range organization. While it cannot be excluded that an intrinsic self-organization of PCP proteins may force the coordination of ciliary beat directions, our observations rather suggest that the organization of PCP proteins is the result of hydrodynamic forces. On a local scale however, one could put forward that before tissue confluence, the intrinsic turbulent dynamics of jamming cells could generate small circular structures^21^ that could facilitate the emergence of local swirls. At large scale, in-vivo, internal forces exerted on the epithelium during embryogenesis are involved to set the planar polarization ^22^. In in-vitro cultures, these forces do not exist and it is the hydrodynamic forces that could lead to the large-scale self-organization of PCP proteins. This would explain the circular organization of PCP and the plasticity of mature cultures allowing the reorganization of the directions of ciliary beats under the effect of hydrodynamic forces.

We conclude with two remarks, First, our model extends the class of Vicsek-like models ^23,24^ based on simple rules between moving individuals and used to describe the collective behavior observed in active biological matter, such as bacterial colonies, cytoskeletal elements and epithelial monolayers ^25–27^. This approach paves the way for the description of novel complex systems made of non-motile active constituents where interactions arise from external forces such as hydrodynamics. Second, from an applied perspective, our results could provide important insights for adapting gene therapy strategies, for example to determine the number of ciliated cells that need to be repaired in primary ciliary dyskinesia.

## Materials and methods

### In-vitro bronchial epithelium

Cultures of human bronchial epithelium reconstituted from primary cells, were either bought from Epithelix (MucilAir) or cultured from human transplant donor lungs deemed unsuitable for transplantation and donated to medical research. The ethics committees of the institutions involved approved this study (CERC-SFCTCV-2018-5-6-9-8-32-DjXa). Primary human bronchial epithelial cells were isolated by protease digestion of human airways, and cells were cultivated under Air-Liquid Interface (ALI) conditions, as previously described ^14,28^. All experiments were performed in accordance with relevant guidelines and regulations.

The tissue is reconstituted at the Air Liquid Interface in transwells of 6 mm in diameter. The apical side of the tissue is at the air interface and the basal side is in contact via a porous membrane (pore size of 0.4 μm) with the culture medium: Epithelix differentiation culture medium during ciliogenesis, then, Epithelix MucilAir culture medium; Pneumacult ALI medium (Stem cell) for homemade cultures. The culture medium (700 μl) is replaced every two days.

#### Removal of mucus

For experiments in absence of mucus, we remove the mucus by performing an apical washing. We add 200 μl of culture medium on the apical side for 10 minutes, followed by three rinses with the culture medium. After an apical washing, a thin layer of culture medium (~ 10 − 40 μm) remains on top of the cilia. Eventually, if patches of mucus remain stuck on the surface of the epithelium we add 10 μl of culture medium on the apical side that we leave overnight.

### Imaging

We imaged the cultures in bright field on an inverted Nikon Eclipse Ti microscope with a x20 objective and a Luminera infinity camera. The temperature is maintained at 37 °C and a humidified air flow with 5% CO2 is applied to maintain a physiological pH. To image the cultures at the same locations on different days as well as to image the samples on large areas (5×5 tiles), we used a motorized stage calibrated in the xy plane and saved the different positions. First, we visualized the streamlines of the mucus flows by following cellular debris trapped in the mucus to characterise the emergence and growth of mucus flow patterns. Then, after slightly diluting the mucus, by adding of 3-4 μl of culture medium on the apical side to get rid of debris, we imaged the underneath beating cilia in the same area. To quantify the ciliogenesis we imaged on the central part of each sample 25 tiles resulting in an area of about 2mm × 1.5mm. For imaging planar polarity of Vangl1, samples were mounted between a glass slide and a coverslip embedded on a drop of ProLong mounting media. Images were taken with the confocal microscope ZEISS LSM 880 reverse standing AxioObserver 7. The objective used were Zeiss 20x Plan Apochromat 20x/0.8. Two-colors confocal z-series images were acquired using sequential laser excitation, and a z-slices interval 1μm, using Tile Scan mode covering areas of ~2×2 mm^2^. Images were converted into single plane projection (MIP) and analyzed or edited using ImageJ software.

### Immunofluorescence staining of Vangl1 and ZO1

Samples were first fixed in 100% methanol, 10 minutes at −20°C. Then, washed 3×3min in PBS at 4°C and blocked with BSA 3% in PBS 30min at RT. Primary antibodies against Z01 (mouse monoclonal IgG1 Thermo Scientifique 33-9100) and Vangl1 (rabbit Sigma Aldrich HPA025235) were diluted at 1/200 and 1/500 respectively in blocking medium and incubated 45min at RT. After 2×5min washes with PBS-Triton 0.1%, secondary antibodies: Alexa mouse IgG1 568 (ThermoFisher A21124) diluted at 1/800 and Alexa Fluor anti-rabbit 488 (Life Technologies A21206) diluted at 1/500 were incubated in blocking medium 30min at RT. Finally, samples are washed 2×5min. The membrane of the Transwell was cut and mounted in the coverslip.

### Quantification of Vangl1 orientation

We measured manually the orientation of Vangl1 protein on each individual cell in the regions of interest using imageJ software. Then we plotted the distributions of beat directions according to the measured Vangl1 orientations with beat direction = Vangl1 orientation + 90°.

### Image processing and quantification

Cilia density *φ*, beating directions of cilia and localisation of topological defects were quantified using in-house image processing programs implemented in python and applied on videomicroscopy acquisitions (40 fps). We first performed a standard deviation projection over 40 frames to reveal the trajectory of the tip of the cilia. We binarised this image by applying a threshold to compute *φ*. We implemented the algorithm described in Püspöki et al. ^29^ based on the structure tensor computation to determine the direction of each pixel on previously obtained standard deviation projection images. The director field of ciliary beats was computed by averaging the orientation of individual pixels in a box of 15×15 pixels, corresponding to the size of a single ciliated cell. Then we localised the topological defects from the director field by computing the winding number^30^.

To visualize the streamlines of the mucus flow on a still image, we perform a temporal standard deviation projection from video acquisition on 100 to 300 frames depending on the frame rate.

### Numerical simulations

Two-dimensional simulations are performed using a D2Q9 lattice-Boltzmann method based on a BGK collision model ^17^. The momentum source modeling the effect of the ciliated epithelium on the flow is taken into account using the forcing model proposed by Guo et al. ^31^. The flow is described on a uniform and cartesian numerical grid. The distance between two neighboring nodes is –, where D denotes the side length of hexagonal elements, i.e. each hexagonal element contains 65 numerical nodes, approximately. At each time step, the flow velocity is averaged on each ciliated elements, allowing to compute the local angle between the ciliary force and flow directions, called on the i-th ciliated element. The orientation of the ciliary forcing is updated following a streamwise-alignment rule, namely

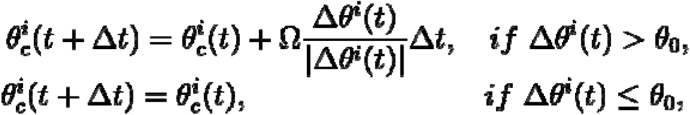

where is the angle between the ciliary force and a given direction on the i-th ciliated element at time t, is the model time step, is a rotation rate and is threshold angle to trigger active realignment. The parameter drives the transient re-reorientation of the ciliated elements but does not affect the final steady solutions presented in this work. It is set to —, where designates the typical mucus velocity. A small threshold angle is employed, namely radians using the present numerical parameters, so that its influence on model solutions can be neglected. The number of performed time steps varies from one computation to the other, since simulations are systematically continued until a steady solution is reached. This is achieved by checking the statistical convergence of several global quantities as the total fluid kinetic energy and the space-averaged instantaneous variation of ciliary-force orientations.

## Data availability

Data that support plots and other findings within this manuscript are available from the corresponding authors upon reasonable request.

## Code availability

Custom codes that were used to analyse experimental data within this manuscript are available from the corresponding author upon reasonable request.

## Acknowledgements

The authors thanks surgeons Pascal Thomas and Xavier Benoit D’Journo, service de chirurgie thoracique, Hôpital NORD, AP-HM, for providing biological material.

## Author contributions

E.L and A.V designed the study. S.G, U.D and J.F conceived the computational model. E.L designed and carried out image analysis. E.L and C.J. acquired experimental data. A.N. acquired the data on PCP. E.L, CJ and A.N analyzed experimental data. S.G. implemented and performed simulations. D.G. performed bronchial epithelium cultures. All authors discussed the results and the manuscript. E.L, S.G and A.V wrote the paper.

## Competing interests

The authors declare no competing interests

