## Supplementary information for "Active mucus-cilia hydrodynamic coupling drives self-organisation of human bronchial epithelium"

| # sample | # patient | origin |
|---|---|---|
| 1 | 1 | Hospital |
| 2 | 2 | Epithelix |
| 3<br>4<br>5 | 3 | Epithelix |
| 6<br>7<br>8 | 4 | Hospital |
| 9<br>10 | 5 | Epithelix |
| 11 | 6 | Hospital |
| 12<br>13 | 4 | Hospital |
| 14<br>15<br>16 | 7 | Hospital |
| 17 | 8 | Epithelix |
| 18 | 9 | Epithelix |
| 19 | 10 | Epithelix |
| 20 | 11 | Hospital |

**Table S1 Samples used in this study**

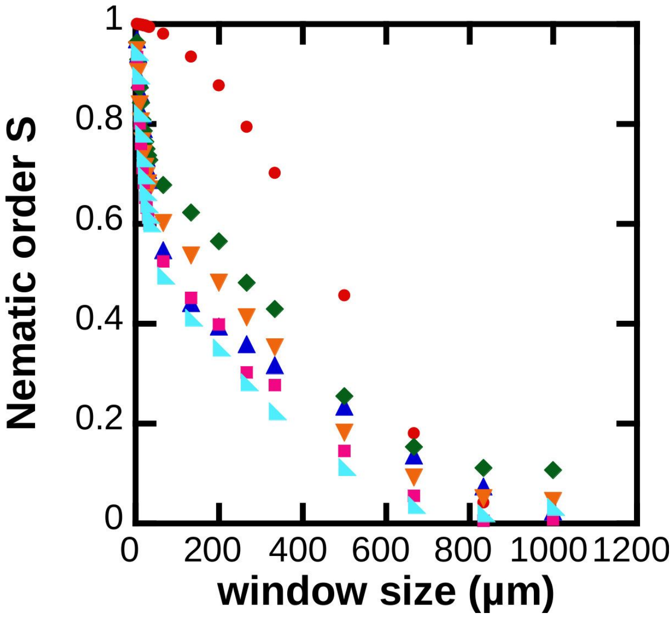


**Figure S1:** Parameter order of the directions of ciliary beats underneath millimetric swirls formed after addition of diluted mucus on disorganised epithelia. Red circle represent the reference of a perfect circular orientation. Blue triangles, sample #18; green diamonds, sample #15; orange down triangles, sample #14; rose squares, sample #16 and turquoise rectangle triangle, sample #17.

**Supplementary Videos:**

**Video 1:** For a cilia density lower than 15% the mucus remains still while the cilia are beating underneath. One can see heterogeneities is the mucus vibrating due to the beating cilia.

**Video 2:** Onset of mucus transport for $\varphi$ ~ 25% on day 12 sample #13.

**Video 3:** Pattern formation on day 13, sample #13, same field of view than movie 2.

**Video 4:** Swirl of mucus on day 14, sample #13, same field of view than movie 2.

**Video 5:** Example of two swirls of mucus which grow by successive accumulation of mucus transported their vicinity. On left side, the swirl is from sample #8, day 12 and on right side the swirl is from sample #7, day 12.

**Video 6:** Millimetric swirl resulting from the growth of the swirl presented on movie 5 (left side), sample #8, day 15.

**Video 7:** Dehydrated mucus gets stuck above the cilia on a mature epithelium.

**Video 8:** After addition of diluted mucus on the surface of a disorganised epithelium, a new millimetric mucus swirl emerges at the epithelial surface, associated with a strong circular orientational order of underlying ciliary beats (Fig. 4c). The field of view correspond to the central part of Fig. 4c.
